# Using the Price equation to detect inclusive fitness in class-structured populations

**DOI:** 10.1101/2020.03.09.982892

**Authors:** António M. M. Rodrigues

## Abstract

Inclusive fitness theory has transformed the study of adaptive evolution since 1964, contributing to significant empirical findings. However, its status as a theory has been challenged by the proposals of several alternative frameworks. Those challenges have been countered by analyses that use the Price equation and the regression method. The Price equation is a universal description of evolutionary change, and the partitioning of the Price equation using the regression method immediately yields Hamilton’s rule, which embodies the main tenets of inclusive fitness. Hamilton’s rule captures the intensity and direction of selection acting on social behaviour and its underlying causal structure. Recent work, however, has suggested that there is an anomaly in this approach: in some cases, the regression method fails to estimate the correct values of the variables in Hamilton’s rule and the causal structure of the behaviour. Here, I address this apparent anomaly. I argue that the failure of the simple regression method occurs because social players vary in baseline fecundity. I reformulate the Price equation and regression method to recover Hamilton’s rule and I show that the method correctly estimates its key variables. I show that games where baseline fecundity varies among individuals represent a more general set of games that unfold in class-structured populations. This framework supports the robustness and validity of inclusive fitness.

## Introduction

Inclusive fitness (Hamilton 1964b, a) is thought by some (e.g. Davies et al. 2012) to be one of the most significant contributions to evolutionary theory since Darwin’s (1859) work on Natural Selection. It provides the theoretical foundations for topics that range from sex allocation (Charnov 1982, West 2010) and the evolution of altruism (Bourke 2011) to parent-offspring conflict (Trivers 1974, Haig 2002) and dispersal evolution (Hamilton and May 1977, Clobert et al. 2012), and it contributes to our understanding of major evolutionary transitions in individuality (Maynard Smith and Szathmáry 1995, Boomsma 2009, Bourke 2011). Despite its explanatory power, inclusive fitness is a concept that has also been the subject of a good deal of controversy. Some argue that inclusive fitness fails when games deviate from additivity (e.g. van Veelen 2009); others claim that it cannot fully explain group selection and that it requires weak selection or rare mutants (e.g. Wilson and Wilson 2007, van Veelen 2009); and still others suggest that it fails to provide a causal account of social behaviour and cannot be empirically tested (Allen et al. 2013, Nowak et al. 2017).

The Price equation has been the main mathematical tool used to address these critiques of inclusive fitness (Queller 1992b, Gardner et al. 2011). It is a universal description of evolutionary change (Price 1970, 1972, Hamilton 1975, Frank 1997, Queller 2017) that supports the analysis of evolutionary quantitative genetics (Lande and Arnold 1983), indirect genetic effects (Moore et al. 1997), and multi-level selection (Okasha 2006). That the Price equation provides a framework for inclusive fitness was first proposed by Hamilton (1970). It has been developed by many since then (Grafen 1985, Queller 1992a, b, Frank 1997, Grafen 2000, Gardner 2015, Grafen 2015), including those who deploy it to address critiques (Queller 1992b, a, Gardner et al. 2011, Rousset 2015). It defines fitness costs and benefits as partial regression coefficients that emerge from an analysis of social behaviour (Queller 1992b, a, Gardner et al. 2011, Rousset 2015). The regression approach has been suggested to demonstrate that inclusive fitness is as general as natural selection and that the actor-centric interpretation of behaviour remains the most robust paradigm in social evolution, both from the theoretical and empirical standpoints (Gardner et al. 2011, West and Gardner 2013).

This view of social evolution, however, has been challenged. In particular, Allen et al. (2013) and Nowak et al. (2017) identified a set of games where the regression analysis fails to yield the correct values of the costs and benefits of the games’ social interactions. This failure of the regression approach called into question the logical status of inclusive fitness within evolutionary biology, in particular raising the issue of whether inclusive fitness can in principle provide a correct account of social behaviour (Birch 2014, Birch and Okasha 2015, Akçay and Van Cleve 2016, Okasha 2016). Some are now starting to question whether inclusive fitness provides a solid framework for the development of novel hypotheses, the design of experiments, and the interpretation of empirical data (e.g. Gadagkar 2016, Whiteley et al. 2017).

It is thus crucial to understand why specific types of games cause the regressions used in inclusive fitness models to break down. Here, I ague that variation in the baseline fecundity of social partners is the underlying cause of the failure of the simple regression method. Understanding this class of games requires an extended version of the Price equation and the regression method. I show that the extended version of the Price equation recovers a form of Hamilton’s rule that while not exactly identical to Hamilton’s original formulation it follows the same logic. I then show that the games in which individuals vary in baseline fecundity belongs to a wider set of class-structured games with broad empirical significance.

## The Price equation

The Price equation is a mathematical statement about how properties of a population of entities change over time (Price 1970, 1972). More precisely, it maps the relationship between two sets of entities and it describes how average quantities change from one set to the other (Frank 2012). Typically, one set is called the parental generation and the other the offspring generation. The entities of these two sets are connected by directed acyclic graphs that define multiple family trees, where the source nodes are the entities in the parental population and the outgoing nodes are the entities in the offspring population (Fig. 1A; Gardner 2020). The entities of the sets are assumed to vary in their breeding value, which can be inherited from parents to offspring with different degrees of fidelity. These assumptions, depicted in diagram 1A, give rise to the Price equation, which describes changes in the breeding value that occur between the parental and offspring population (see Gardner 2008, Frank 2012 for reviews, and the appendix for details). Changes in mean breeding value can occur for two main reasons: natural selection and transmission biases (Frank 1997, Okasha 2006, Gardner 2008). Here, I focus on changes in breeding value due to the action of natural selection. Under these conditions, the most general form of the Price equation is given by

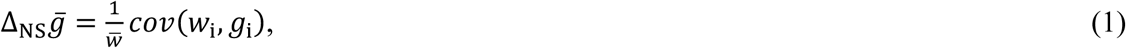

where: *w*_i_ is the reproductive success of the *i*th individual in the population; *g*_i_ is the breeding value of the *i*th individual; 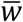 is the average reproductive success in the population; 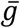 is the average breeding value in the population; and 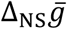 denotes the change in the average breeding value between the parental and offspring generations owing to the action of natural selection. This statement does not depend upon any assumption regarding the nature of the population; it therefore provides a general description of the action of natural selection (Price 1970, Gardner 2008, Frank 2012). The Price equation tells us that the change in the average breeding value between generations is given by the covariance between the relative reproductive success of individuals and their breeding value.

**Figure 1.**
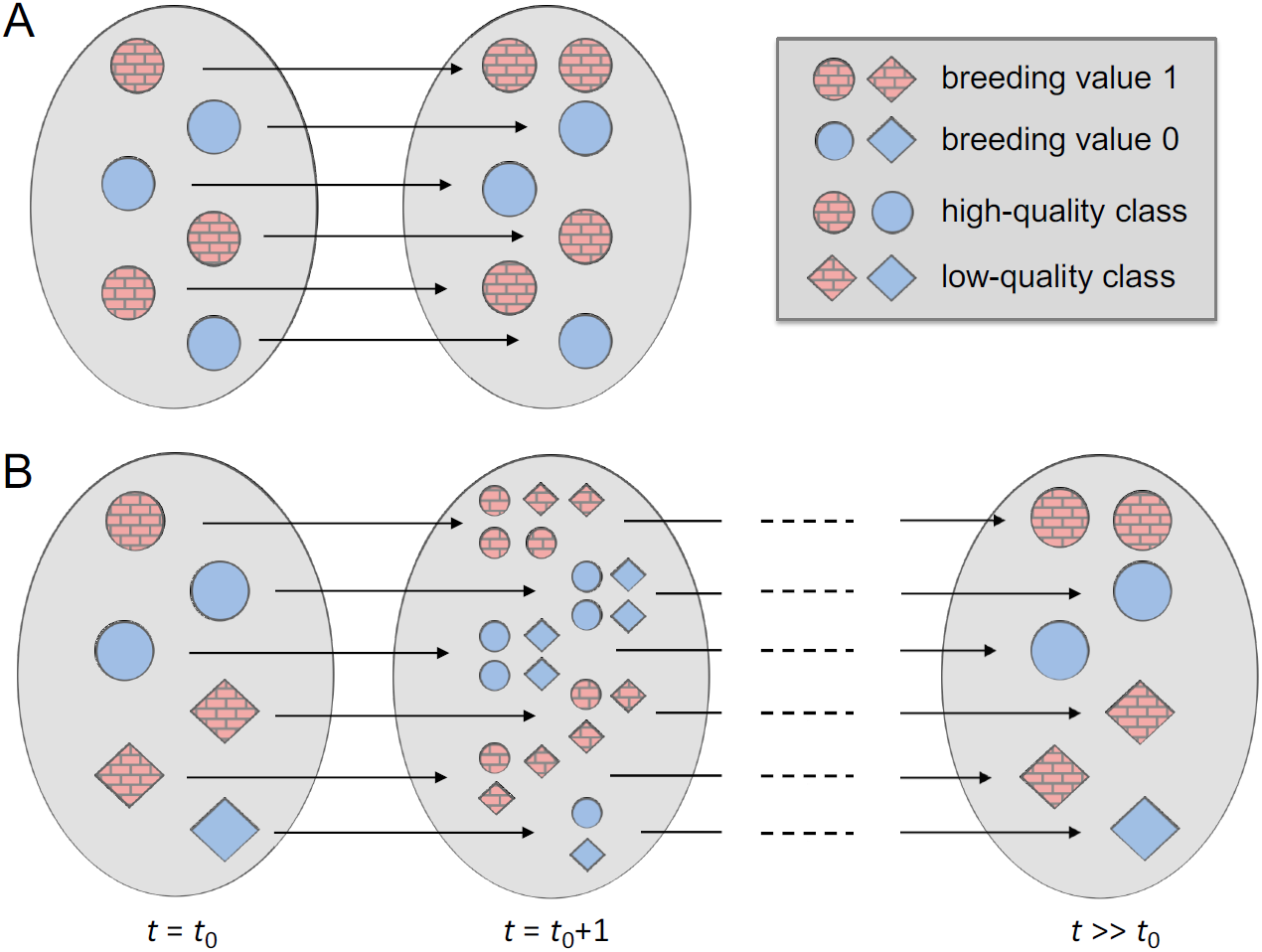
Acyclic direct graph describing the dynamics of the population. The left-most population is the parental population, while the middle and right-most populations are the descendant populations. The colour of each entity represents the breeding value of an individual, while shape represents their class. **A**. Visual depiction of the standard Price equation where no variation in quality is considered. **B**. The dynamics of a population when individuals vary in quality. The left-most panel represents the current population at time *t*_0_, while the second panel represents the population in the next time step (i.e. *t* = *t*_0_ + 1). The right-most panel represents a descendant population in the distant future (i.e. *t* >> *t*_0_).

## The Price equation extended

### The Price equation in a class-structured world

The standard derivation of the Price equation assumes that all entities in the population are identical except for their breeding value, as represented in diagram 1A (e.g. Price 1970, Gardner 2008, Frank 2012). Conceptually, we can modify this framework in three main ways. First, rather than two sets of entities, the parental and offspring populations, we can consider more than two sets of populations. For instance, we can imagine that entities in the first population give origin to entities in the second population, entities in the second population give origin to entities in the third population, and so forth. Second, rather than undifferentiated individuals (or entities), we can consider that individuals differ in a property, which we can call quality, and which we represent by different shapes in the diagram 1A. Third, we can allow the quality of individuals to influence both the number of entities they produce, as well as the quality (or class) of the entities they produce, where quality is any phenotype of an individual that affects its fitness (see diagram 1A). Although quality often defines classes (e.g. large and small individuals), classes exist even if there are no obvious phenotypic differences among individuals, such as when individuals occupy habitats of different quality (e.g. core and marginal habitats).

The aim is to discover how the average breeding value of a population in the future is affected by natural selection acting on the current generation. To do so, we partition total fitness into a current and future component. Current fitness, denoted by *w*_ij→1_, is the contribution of the *i*th individual in the current population to the offspring population, where *j* is the class of the focal individual and *l* is the class the individuals produced by the *i*th individual. Future fitness, denoted by *v*_1_, is the contribution of a class-*l* individual in the offspring generation to a population in the future. Future fitness, or reproductive value, is calculated using the “counter-factual” method by considering a neutral population from time *t*_0_ + 1 onwards (Frank 1998, Gardner 2015). This enables us to differentiate natural selection acting on the current generation from natural selection acting on subsequent generations (Frank 1998, Gardner 2015).

As in the previous section, the assumptions underlying diagram 1B give rise to a corresponding “Price equation” (see appendix for details), which is given by

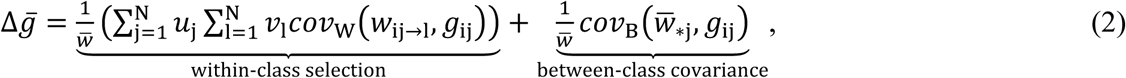

where: *N* is the different classes (or qualities) of individuals in the population; *u*_j_ is the frequency of individuals in class-*j*; *g*_ij_ is the breeding value of the *i*th individual in class-*j*; and 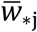 is the mean fitness of individuals in class-*j*. I use *cov*_W_ and *var*_W_ to denote covariances and variances within any given class, and *cov*_B_ and *var*_B_ when covariances and variances are taken between classes and across all individuals in the population.

This formulation of the Price equation isolates two key processes driving evolutionary change. First, the “within-class selection” terms describes statistical associations between breeding value and fitness within each class, with each covariance being weighted by the frequency of individuals within each class and by the reproductive values of the recipient classes. Note that breeding values may be positively associated with fitness in some classes, but negatively associated with fitness in others. The overall effect depends both on the strength of each association and on the frequency of individuals in each class and on the reproductive values of the recipient classes. The covariance terms within each class can be written as 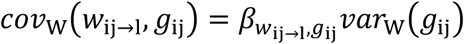. That is, for selection to operate within each class, there must be genetic variation within that class (i.e. *var*_W_(*g*_ij_) > 0) and there must be a statistical association between breeding value and fitness 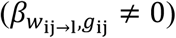. If either of these conditions are not met, then there is no scope for selection to act within that class.

Second, the last term represents selection that operates between classes and / or class effects, which is given by the covariance between breeding value and the mean fitness of each class. If the between-class covariance occurs because of gene action, we call it “selection between classes”. Otherwise, we call it “class-effects”. The covariance between classes is positive whenever higher values of breeding value are statistically associated with higher values of class mean fitness (i.e. higher 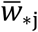), and negative whenever higher values of breeding value are statistically associated with lower values of class mean fitness (i.e. lower 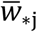). If individuals are randomly distributed across the different classes, then there is no statistical association between breeding value and class mean fitness. In that scenario, the selection between classes and / or class-effects are zero (i.e. 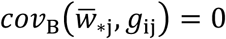), and selection within classes is the only force governing change in average breeding value.

### Classes and the regression approach

In the previous section, I did not specify the relationship between fitness (or reproductive value) and breeding value. In the context of kin selection, the fitness of a focal individual will depend both on its own breeding value and on the breeding value of its partners. This relationship between reproductive success (the dependent variable) and breeding values (the independent or predictor variables) can be described by a statistical model as part of a regression analysis (Queller 1992b, a).

The form of the statistical model depends on the covariance expressions in the Price equation. Covariances in the first term of the Price equation are calculated across the set of individuals within each class, while the covariance in the second term is calculated across the set of all individuals in the population. Therefore, the regression analysis is performed within each class, when considering the first (within-class selection) term, but across all individuals, when considering the second (between-class covariance) term.

#### Within-class selection

Let us start by focusing on the regression analysis within each class. For each class, I denote the intercept of the statistical model by *β*_0j_, where *j* represents the focal class. In addition, the fitness of a focal individual in class-*j* depends on the breeding value of the focal individual, on the breeding value of the individuals in the same class, and on the breeding value of individuals in other classes. Thus, the estimated fitness of the focal *i*th individual in class-*j* can be written as

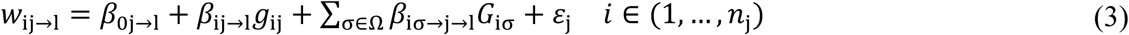

where: *β*_ij→1_ is the partial regression coefficient that gives the effect of the focal individual’s breeding value on its own fitness when the focal individual produces class-*l* individuals; *β*_iσ→j→1_ is the partial regression coefficient that gives the effect of a class-*σ* social partner on the fitness of the focal class-*j* individual when the focal individual produces class-*l* individuals; *g*_ij_ is the breeding value of the focal individual; *G*_iσ_ is the breeding value of the focal individual’s class-*σ* social partners; *n*_j_ is the number of individuals in class-*j*; and, finally, *ε*_j_ is the uncorrelated error between the observed and estimated values.

#### Between-class covariance

I now focus on the “between-class covariance” term in the Price equation (equation (2)). Let each class be defined by its mean fitness 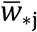, and denote *σ*_ij_ as the class phenotype, which is defined in relation to the mean fitness of class-*j*. Specifically, I define the class phenotype of the *i*th individual in class-*j* as 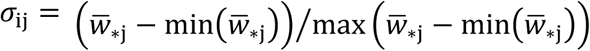, such that the class phenotype *σ*_ij_ is bounded between 0 and 1. This will not affect the calculations because I am simply rescaling the mean fitness of the class. The mean fitness of an individual in a class-*j* can then be described by the following model

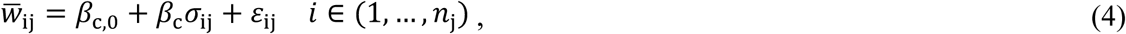

where *β*_c,0_ is the intercept, and *β*_c_ is the effect of the class phenotype on mean fitness. I can now replace this equation in the “between-class covariance” term in the Price equation (equation (2)) to obtain

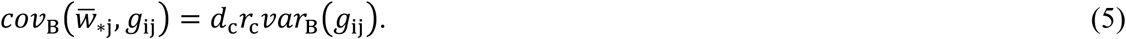

where *r*_c_ = *cov*_B_(*σ*_ij_, *g*_ij_)/*var*_B_(*g*_ij_) is the regression of breeding value on class phenotype, *d*_c_ = *β*_c_ is the effect of class phenotype on mean class fitness. The regression of breeding value on class phenotype, *r*_c_, can be seen as a “class coefficient” that contains information about how breeding value is spread across the different classes. The right-hand side of equation (5) has a pleasant interpretation. The partial coefficient of correlation *β*_c_ gives the effect of class phenotype on the mean fitness of an individual; the class coefficient *r*_c_ gives the association between breeding value and class; and *var*_B_(*g*_ij_) gives the additive genetic variance in the population. We can now pinpoint the conditions under which the covariance between classes (i.e. selection between classes and / or class-effects) is zero. First, the covariance between classes is zero when the genotypes are uniformly distributed among all classes, and therefore when the mutant and neutral allele occur in the same proportions within each class (i.e. *r*_c_ = 0). Second, the covariance between classes also vanish when class does not affect mean fitness (i.e. *d*_c_ = 0). Third, the covariance between classes is zero in the absence of additive genetic variance in the population (i.e. *var*_B_(*g*_ij_) = 0).

## Hamilton’s rule in a class-structured world

From the Price equation and the regression analysis, Hamilton’s rule for different forms of social behaviour can be derived. Here, I will focus on two forms of behaviours: first, behaviour that affects the fecundity of both actors and recipients (fecundity effects); second, behaviour that affects the survival of both actors and recipients (survival effects).

I start with a general model for the fitness of a focal individual and allow it to derive fitness from the production of offspring and from its own survival. I define the class-specific fitness of a focal individual as

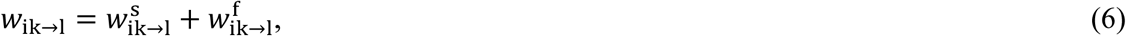

where 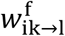 is the fecundity component, and 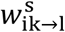 is the survival component of fitness.

### Fecundity effects

When focusing on fecundity alone, I assume that there is standing additive genetic variance for fecundity but not for survival. Because fecundity is the trait of interest, I need to define how fecundity influences the overall reproductive success of a focal individual. Let the reproductive success of the *i*th individual in class-*k* through offspring that become class-*l* individuals be given by 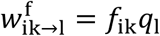, where *f*_ik_ is the fecundity of the *i*th class-*k* individual and *q*_1_ is the fraction of rank-*l* offspring produced by a focal mother. Here I assume that mothers vary in their fecundity, but they produce the same proportions of the different types of offspring.

I now need to define how social interactions unfold. Let actors belong to class-*α*, and recipients belong to class-*ρ*, with ρ ∈ Θ, where Θ is the class of all recipients. From equation (2), I obtain

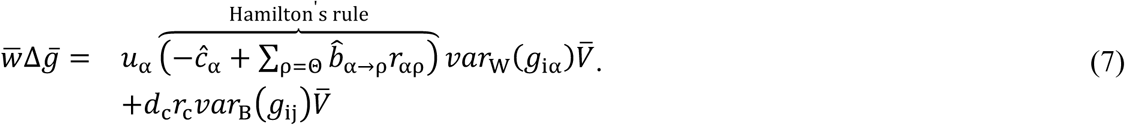

where: 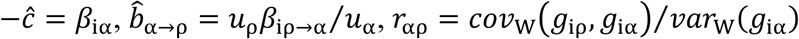, and 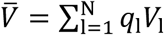 is the expected reproductive value of offspring (see Appendix for details). Note that the only assumptions are that additive genetic variation affects fecundity alone, and that there is no transmission of class from parents to offspring. The interpretation of this form of Hamilton’s rule is straightforward, closely following the canonical interpretation. The focal actor pays a cost *c*_α_ to provide a benefit to a set of recipients Θ. Each recipient enjoys a benefit *b*_α→ρ_, which must be depreciated by the coefficient of relatedness *r*_αρ_ between actor and recipient.

### Survival effects

Now consider survival effects. Here I assume that there is standing genetic variation for survival, but not fecundity, and therefore the fecundity component of fitness does not affect our calculations. The fitness of a mother can be written as 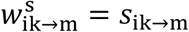, where *s*_ik→m_ is a mother’s survival probability. Performing the regression analysis outlined in the preceding section, I find that the mean change in breeding value due to the action of natural selection becomes

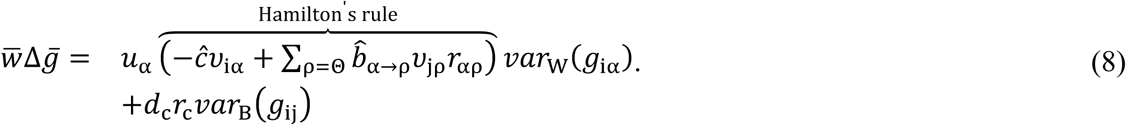

where: *υ*_iα_ is the future reproductive value of the actor, and *υ*_jρ_ is the future reproductive value of recipients. Thus, under survival effects, I find that the estimated costs and benefits must be weighted by the expected reproductive value of actor and recipients, respectively. Here the little *c*’s and *b*’s denote short-term costs and benefits, with reproductive value converting short-term costs and benefits into long-term fitness effects. Nevertheless, the general form of Hamilton’s rule remains identical to more standard forms of Hamilton’s rule.

## Detecting inclusive fitness

Let me now illustrate how this framework can be used to analyse and understand concrete evolutionary games. In particular, I will employ the framework to analyse the examples used by Allen et al. (2013) and Nowak et al. (2017) to identify several types of evolutionary games in which the simple Price equation-regression approach to social evolution breaks down. I then discuss examples that explicitly contrast the simple regression analysis with one enhanced by the class-structured form of the Price equation. I first consider a game where individuals associate with each other but no real social transactions occur (cf. Fig. 1 and Fig. 2A in Allen et al. 2013). Next, I consider a game in which high-fecundity individuals help low-fecundity individuals (cf. Fig. 2C in Allen et al. 2013). Then, I consider a game in which low-fecundity individuals inflict a cost on high-fecundity individuals (cf. Fig. 2B in Allen et al. 2013). I will focus on selection between consecutive generations. Further, I assume that the between-class covariance is not due to the action of genes, and therefore I will use the term “class-effects” to refer to this covariance. I will return to this subject below.

**Figure 2.**
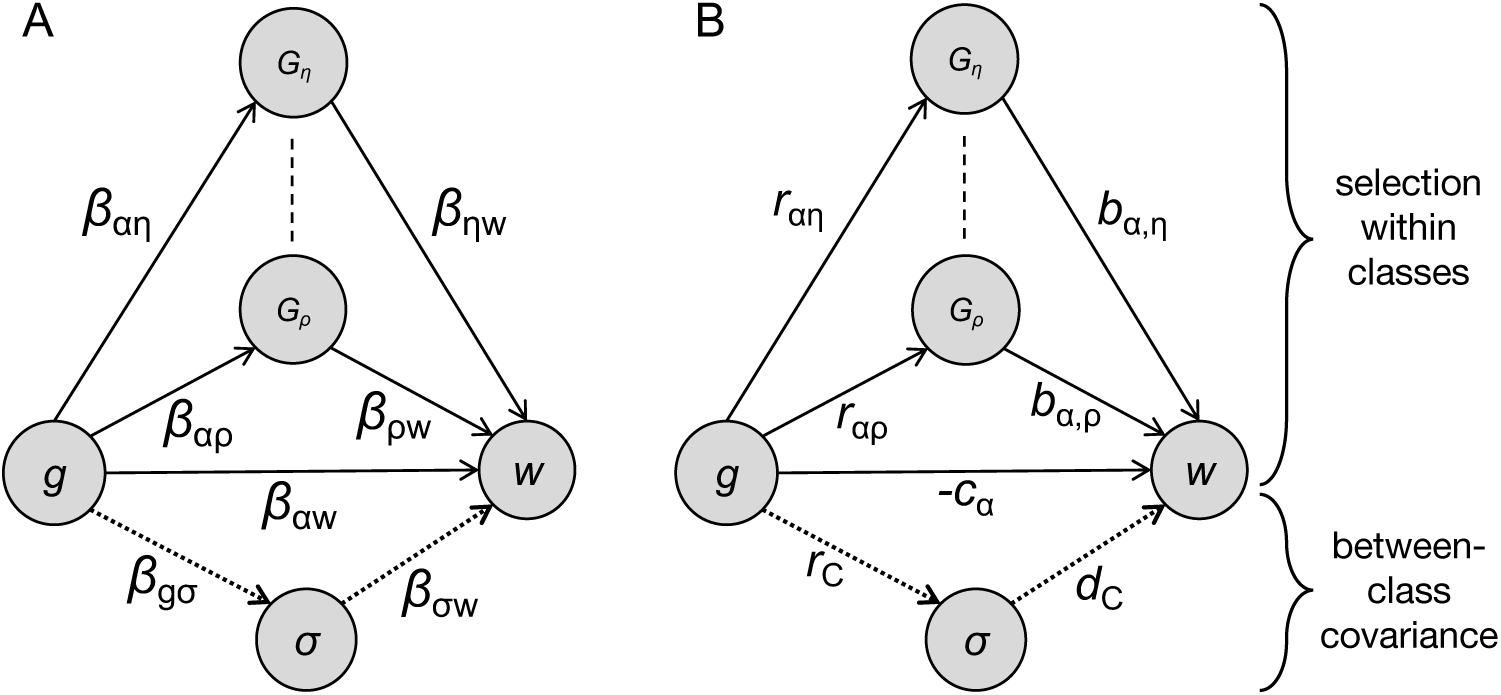
Path diagram with the causal model describing the association between breeding value and fitness. Fitness (*w*) depends on the breeding value of the focal individuals (*g*), on the breeding value of the focal’s social partners (*G*) and on class phenotype (*σ*). **A**. Each edge is weighted by a partial coefficient of correlation. **B**. Each edge corresponds to a variable in Hamilton’s rule. For instance, the direct association between fitness and breeding value is the additive inverse of the behaviour’s cost (– *c*), the association between breeding values gives the relatedness coefficient (*r*), and the association between breeding value and class phenotype gives the class coefficient (*r*_c_).

### Anomalies in previous literature

Most of the anomalies identified by Allen et al. (2013) and Nowak et al. (2017) occur because they did not take into account the underlying class-structure of the games. When a population has class-structure, gene frequency change can occur because of within-class selection or because of a nonzero between-class covariance (due to either selection or class-effects). If one does not properly represent the classes in the Price equation, then selection is compressed into a single regression coefficient that includes both selection within classes and the covariance between classes; that move affects the estimates of costs and benefits of behaviours.

Let us consider the game provided in Fig. 1 in Allen et al. (2013). First, because individuals differ in their baseline fitness, which can take the values 4, 2 and 0, class must be taken into account. Second, because one class is composed of a single individual – i.e. there is a single individual with baseline fitness 4 – there is no scope for selection to operate within that class. Third, because the class of individuals with baseline 2 is composed of genetically identical individuals, there is no scope for selection to operate within that class as well. Fourth, while there is scope for selection within the class of individuals with baseline 0, the regression of breeding value on fitness is zero, and therefore selection within the class of individuals with baseline 0 is null as well. Thus, all change in gene frequency must occur because of a nonzero covariance between classes (i.e. class-effects). If our framework is correct, class-effects, as given by equation (5), must be equal to total selection, as given by the standard Price equation in equation (1). That is, 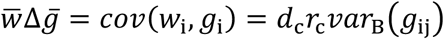. Indeed, we find that *cov*(*w*_i_, *g*_i_) = *d*_c_*r*_c_*var*_B_(*g*_ij_) = 0.125 as expected. Here, we find a nonzero covariance between classes because of a positive effect of class phenotype on baseline fitness (*d*_c_ = 4) and because of a positive association between breeding value and class phenotype (*r*_c_ = 0.133). These issues apply to the example of Fig. 2A in Allen et al. (2013), where there is no selection within classes, either because classes contain a single individual, because classes do not have genetic variation, or because there is no correlation between breeding value and fitness.

In the examples provided in Fig. 2B and 2C there is both selection within classes and a nonzero covariance between classes. In Fig. 2B, there is no selection within the classes with baseline fitness 0 and 5 because they lack genetic variation. However, there is selection within the class composed of individuals with baseline fitness 1, where the regression analysis within that class provides the correct estimate of the cost of the behaviour (i.e. *ĉ* = 1), given by the regression of breeding value on class-specific fitness, as defined above. Selection within that class, however, only captures a fraction of the total selection. The other fraction is given by class-effects, which is *d*_c_*r*_c_*var*_B_(*g*_ij_) = −0.025. Our calculations correctly recover total selection, as given by the standard Price equation (i.e. *cov*(*w*_i_, *g*_i_) = −0.125), for when we add together selection within classes and class-effects, we obtain −0.100 − 0.025 = −0.125, as expected.

Let us consider the four examples given in Fig. 3 in Nowak et al. (2017). In all four, the simple method fails because of class structure. In the example of Fig. 3A, there are three classes: (1) “blue” individuals that interact with other blue individuals; (2) “blue” individuals that interact with “red” individuals; and (3) “red” individuals that interact with “red” individuals. Because there is no genetic variation within any of these classes, there is no selection within classes, and all evolutionary change results from a nonzero covariance between classes. In the examples of Figs. 4B-4D, there are two classes defined by the baseline fitness of individuals. In all three cases, there is again no scope for selection within classes, as classes have no genetic variation, and all evolutionary change is due to a nonzero covariance between classes.

**Figure 3.**
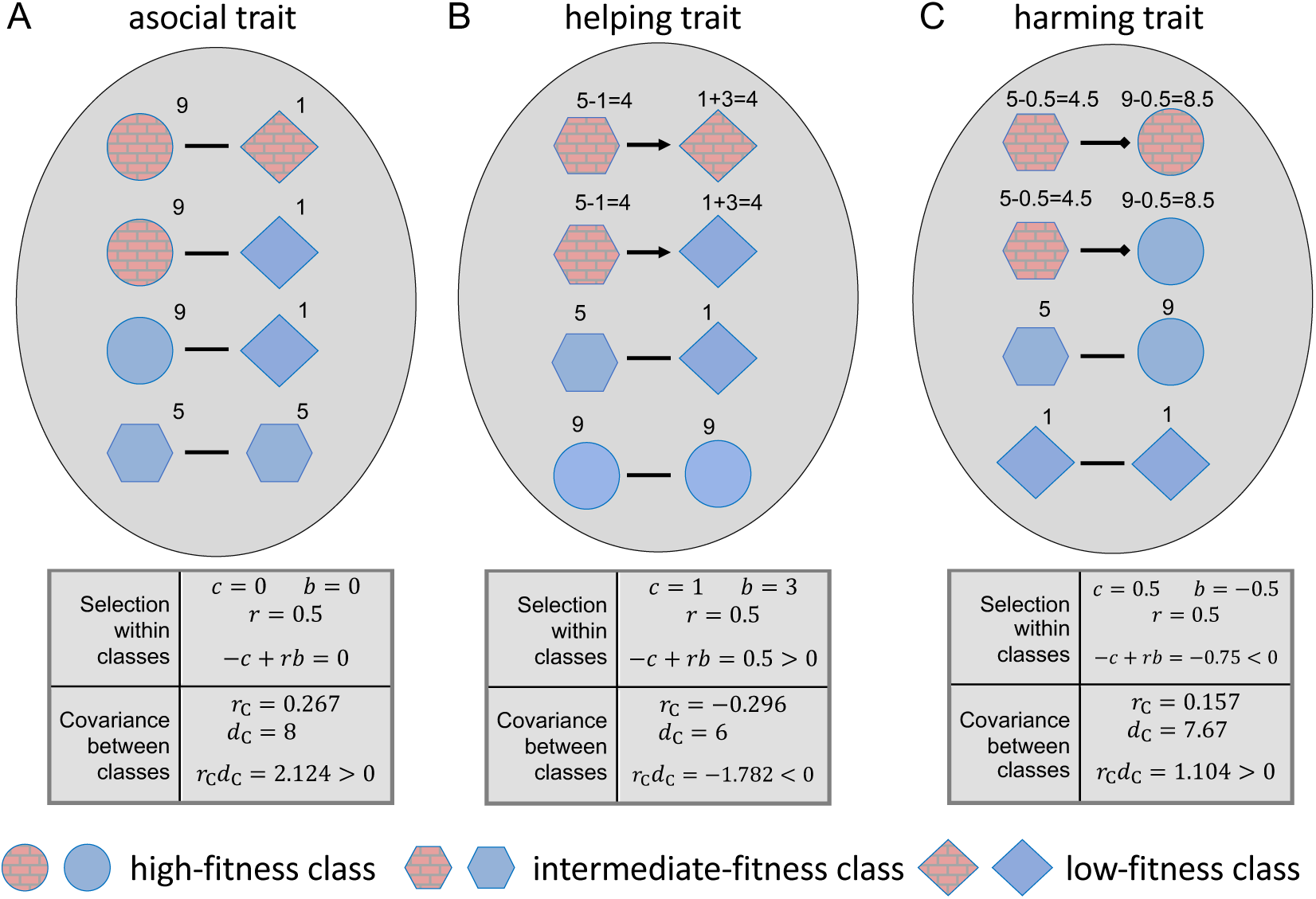
Representation of each evolutionary game. Colour represents breeding value, shape represents baseline fecundity, and numbers represent baseline fecundity with the corresponding increments or decrements owing to social interactions. **A**. Individuals associate with each other but no actual social transactions occur. **B**. Intermediate-fecundity individuals help low-fecundity individuals. **C**. Low-fecundity individuals harm high-fecundity individuals.

**Figure 4.**
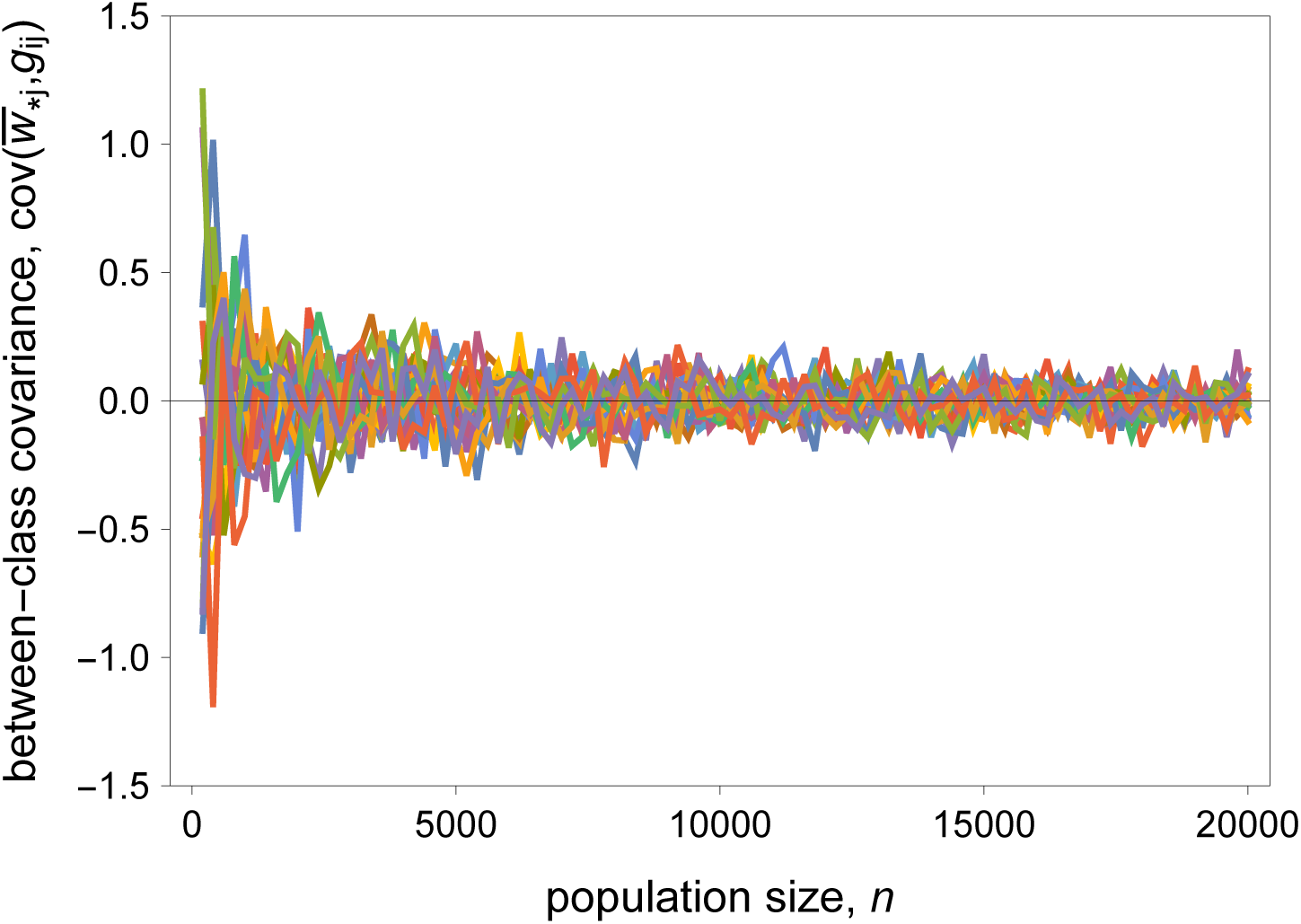
Between-class covariance as a function of population size for different replicates. If population size is relatively small, sampling biases will cause some genotypes to occur at higher frequency in one of the classes. Sampling biases generate a correlation between breeding value and mean fitness. When the population is relatively large, however, sampling biases will become less prominent, and the covariance between breeding value and mean fitness tends to vanish.

### Further examples

Here, I consider cases in which there is scope for selection within classes and a nonzero covariance between classes.

#### No transactions between individuals

Let us consider a game whereby low-fecundity individuals tend to associate with high-fecundity social partners, but no social transactions occur (Fig. 3A; cf. the Hanger-On game in Allen et al. 2013). In other words, social interactions between social partners carry neither costs (*c* = 0) nor benefits (i.e. *b* = 0). I first estimate costs and benefits using the simple regression method. I find that the simple method leads to the wrong estimation of costs and benefits. Specifically, it estimates a negative cost (i.e. *ĉ* = – 4.0) and a negative benefit 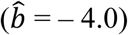, and therefore it incorrectly classifies the behaviour as a selfish trait, when the behaviour is asocial (i.e. *c* = 0 and *b* = 0).

Now I estimate costs and benefits using the regression analysis based on the extended Price equation. I find that the extended regression method correctly estimates the costs and benefits of the social behaviour (i.e. *ĉ* = 0 and 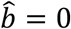). The extended version of the Price equation also explains why the simple regression method fails: it detects correlations between breeding value and class (i.e. *r*_c_ = 0.267) and between class and fitness (i.e. 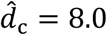). This is because individuals with higher breeding value have an above-average tendency to be in classes of higher fitness, and therefore there is either selection between classes or class-effects. Note that both the simple and the extended regression method correctly predict the intensity and direction of evolutionary change (i.e. 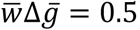), but only the class-based regression method correctly explains the causes of the behaviour.

#### High-fecundity helpers

Here I consider a game in which high-fecundity individuals form one class, and low-fecundity individuals form another, and I assume that high-fecundity individuals help low-fecundity individuals (Fig. 3B; cf. Fig. 2C in Allen et al. 2013). I assume that the cost of the behaviour is one (i.e. *c* = 1) and the benefit is three (i.e. *b* = 3; Fig. 3B). Thus, because both the cost and benefit are positive (i.e. *c* > 0 and *b* > 0), the behaviour should be classified as altruistic. I find that the simple regression method incorrectly estimates costs and benefits: it estimates a positive and incorrect cost (i.e. *ĉ* = 1.091) and a negative and incorrect benefit (i.e. 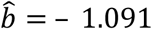). Thus, the simple method incorrectly classifies an altruistic behaviour as spiteful.

In contrast, the regression method based on the class-structured Price equation accurately describes the behaviour: it correctly estimates the costs and benefits of the social behaviour (i.e. *ĉ* = 1 and 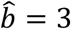, and it explains why the simple regression method fails, for it detects correlations between breeding value and class (i.e. *r*_c_ = −0.296) and between class and mean fitness (i.e. 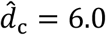). That is, individuals with higher breeding value have a tendency to be in classes of lower mean fitness. As before, both methods correctly predict the direction and intensity of evolutionary change 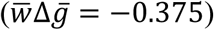, but only the extended method generates the correct causal model for the evolution of the behaviour.

#### Harm by low-fecundity individuals

Now consider a game in which a low-fecundity individual inflicts a cost on a high-fitness social partner at a cost to itself (Fig. 3C; cf. Fig. 2B in Allen et al. 2013). I assume that the behaviour entails a cost of 0.5 (i.e. *c* = 0.5), and a benefit of – 0.5 (i.e. *b* = – 0.5). Because the cost is positive but the benefit is negative, the behaviour is classified as spiteful. Here the simple regression method incorrectly estimates the costs and benefits (*ĉ* = 0.636 and 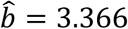). Because both the cost and benefit are positive, the model incorrectly classifies a spiteful behaviour as altruistic.

Again, the extended method yields the correct explanation of the behaviour, for it correctly estimates the costs and benefits of the behaviour (*ĉ* = 0.5 and 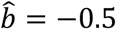) and correctly classifies the behaviour as spiteful. It also clarifies why the simple method fails by detecting correlations between breeding value and class (i.e. *r*_c_ = 0.157) and between class membership and mean fitness (i.e. 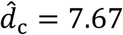). As in the previous examples, both methods correctly predict the selection differential (i.e. 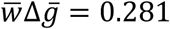), but only the extended method correctly explains the causal reasons underlying changes in gene frequency.

## A closer look at class-effects

Above, we saw that the class-based regression method explains why the simple regression method fails in previous literature and in each of the three examples. In all cases there is a nonzero covariance between classes (either selection or class-effects). That is, there is a correlation between breeding value and class membership and between class and mean fitness. The correlation between breeding value and class is a confounding factor when one uses the simple regression method to estimate costs and benefits, which breaks down as a result. The class-based Price equation captures this “confounding” factor. The confounding factor may be a real biological phenomenon, or an artefact of artificial datasets used to illustrate a hypothetical game. If one specifies that low-fecundity individuals help high-fecundity individuals, then one ought to take into account the distribution of co-operator and defector genotypes among the different classes. If one does not, then one is implicitly assuming that resident and mutant alleles are identically distributed across the different classes. However, the datasets presented above do not fulfil this assumption. For instance, if the dataset is generated at random, then the size of the population and the number of replicates will influence the distribution of genotypes among the different classes. If the probabilities of being a high- or low-fecundity individual are both ½, irrespective of their breeding value, then certain genotypes can be over-represented in high-fecundity classes when the population size is small.

We can illustrate this point by generating random datasets as a function of population size (see Fig. 4). As anticipated, I find that as the size of the population increases, the class coefficient tends to zero (*r*_c_ → 0), and therefore class-effects vanish (Fig. 4). This is because if a population is sufficiently large, the wild-type and mutant allele tend to become equally distributed among the different classes. In contrast, small population sizes contain sampling biases, in which the proportions of wild-type and mutant alleles in each class are not balanced. Alternatively, if the population size is small, but we simulate a sufficiently large number of replicates, the cumulative effect of selection among classes also vanishes (Fig. 4). Note that the data sets used in Allen et al. (2013) and Nowak et al. (2017) contain precisely such sampling bias.

## The elements of the Price equation

Each element of the Price equation provides a description of the different processes that contribute to change in average gene frequency. The frequency of individuals in each class measures the impact of each environment on the intensity of selection. This occurs, for instance, whenever habitats are subdivided into different types. All else being equal, marginal environments (sinks), in which individuals occur at lower frequencies, contribute less to selection than core environment (sources), in which individuals occur at higher frequencies. Thus, the frequency of individuals in each environment is crucial when measuring the influence of each habitat on selection, a classical result (Pulliam 1988). Reproductive value converts current selective pressures into long-term evolutionary change, for an individual in a high-fitness class leaves more descendants than average, and therefore high-fitness individuals are the ancestors of a disproportional number of individuals in future populations. In contrast, individuals that leave no descendants do not contribute to selection through direct reproduction and therefore their reproductive value is zero. The covariances within each class provide a mechanism to standardise the effect of breeding value on fitness by removing class-effects. Variation in weight, size, or body fat, for instance, may be due to environmental factors, rather than the action of genes. The class-specific regression analysis ensures that these environmental effects are stripped away from the changes that are due to the action of natural selection. And the last term in the Price equation captures the statistical association between breeding value and class. This effectively separates class-effects from selection within classes (including kin selection), which is captured by the covariances within each class.

## Further considerations

In the examples outlined above, I have considered games where individuals vary in their baseline fecundity and where social interactions affect the fecundity of actor and recipient. I showed that as long as baseline fecundity is not transmitted from parents to offspring, the reproductive value of offspring can be neglected in Hamilton’s rule, as only the correlations between maternal fecundity and breeding value affect the direction of selection acting on social behaviour. In that scenario, Hamilton’s rule assumes its standard form (Hamilton 1963, Charnov 1977), where the key quantities are the costs and benefits of the social behaviour and the relatedness of the actors and recipients.

In other types of games, for example where survival may vary with class, the reproductive values of actors and recipients must be taken into account. In such cases, Hamilton’s rule deviates from its more common form, in which the costs and benefits of the social behaviour must be weighted by the future reproductive value of actor and recipient, respectively (e.g. Rodrigues 2018). This was foreshadow by Hamilton in his use of *life-for-life* coefficients of relatedness, which include reproductive values (Hamilton 1972). More generally, the approach developed here can be applied to many other types of behaviour, including those in which there are correlations between maternal and offspring quality.

It is important in evolutionary genetics to separate changes in gene frequency ascribed to natural selection from changes in gene frequency that are not due to the action of genes. Fisher pioneered this approach by developing mathematics of gene frequency change that correct for non-adaptive effects (Fisher 1930). Reproductive value and class frequency are crucial concepts in the mathematics of adaptive gene frequency change (Fisher 1930, Taylor 1990, Taylor and Frank 1996, Grafen 2006). The Price equation derived above follows the same principles. Each element of the Price equation corrects gene frequency changes for non-adaptive processes.

In the illustrative examples, I defined classes according to baseline fecundity. More generally, classes can be defined by any phenotypic, behavioural, or social marker (Rodrigues and Gardner 2013). For instance, we may need to classify individuals according to their size, large and small, and their social status, dominant or subordinate. The structure of the population may often require the classification of individuals along multiple dimensions, such as size, age, and social status.

As discussed above, reproductive value converts current selective pressures into long-term adaptive changes (Fisher 1930, Taylor 1990, Grafen 2006, Gardner 2015). But if we are only interested in short-term evolutionary changes, then we simply set reproductive values to one, and the contribution to the offspring population is directly given either by the fecundity or survival of individuals in the current generation.

## Conclusion

The Price equation and the regression method developed in this article provide a general framework for analysing social evolution in class-structured populations. This analysis confirms the pivotal role that Hamilton’s rule plays in explaining social behaviour. The conditions stated here for the evolution of a social behaviour can be traced back to Hamilton’s original derivation and his subsequent work on inclusive fitness.

## Acknowledgements

I thank Andy Gardner and Steve Stearns for comments and helpful discussion.

## References

Akçay, E., and J. Van Cleve. 2016. There is no fitness but fitness, and the lineage is its bearer. Philosophical Transactions of the Royal Society B 371:20150085.

Allen, B., M. A. Nowak, and E. O. Wilson. 2013. Limitations of inclusive fitness. Proceedings of the National Academy of Sciences of the USA 110:20135–20139.

Birch, J. 2014. Hamilton’s rule and its discontents. British Journal for the Philosophy of Science 65:381–411.

Birch, J., and S. Okasha. 2015. Kin selection and its critics. Bioscience 65:22–32.

Boomsma, J. J. 2009. Lifetime monogamy and the evolution of eusociality. Philosophical Transactions of the Royal Society B 364:3191–3207.

Bourke, A. F. G. 2011. Principles of Social Evolution. Oxford University Press, Oxford, UK.

Charnov, E. L. 1977. An elementary treatment of the genetical theory of kin-selection. Journal of Theoretical Biology 66:541–550.

Charnov, E. L. 1982. The Theory of Sex Allocation. Princeton University Press, Princeton, N.J.

Clobert, J., M. Baquette, T. G. Benton, and J. M. Bullock. 2012. Dispersal Ecology and Evolution. Oxford University Press, Oxford, UK.

Darwin, C. R. 1859. On the Origin of Species by Means of Natural Selection, or, the Preservation of Favoured Races in the Struggle for Life. John Murray, London, UK.

Davies, N. B., J. R. Krebs, and S. A. West. 2012. An Introduction to Behavioral Ecology. 4th edition. Blackwell, Oxford, UK.

Fisher, R. A. 1930. The Genetical Theory of Natural Selection. Clarendon Press, Oxford, UK.

Frank, S. A. 1997. The Price equation, Fisher’s fundamental theorem, kin selection, and causal analysis. Evolution 51:1712–1729.

Frank, S. A. 1998. Foundations of Social Evolution. Princeton University Press, Princeton, NJ.

Frank, S. A. 2012. Natural selection. IV. The Price equation. Journal of Evolutionary Biology 25:1002–1019.

Gadagkar, R. 2016. Evolution of social behaviour in the primitively eusocial wasp *Ropalidia marginata*: do we need to look beyond kin selection? Philosophical Transactions of the Royal Society B 371:20150094.

Gardner, A. 2008. The Price equation. Current Biology 18:R198–R202.

Gardner, A. 2015. The genetical theory of multilevel selection. Journal of Evolutionary Biology 28:305–319.

Gardner, A. 2020. Price’s equation made clear. Philosophical Transactions of the Royal Society B 375:20190361.

Gardner, A., S. A. West, and G. Wild. 2011. The genetical theory of kin selection. Journal of Evolutionary Biology 24:1020–1043.

Grafen, A. 1985. A geometric view of relatedness. Oxford Surveys in Evolutionary Biology 2:28–90.

Grafen, A. 2000. Developments of the Price equation and natural selection under uncertainty. Proceedings of the Royal Society B 267:1223–1227.

Grafen, A. 2006. A theory of Fisher’s reproductive value. Journal of Mathematical Biology 53:15–60.

Grafen, A. 2015. Biological fitness and the Price Equation in class-structured populations. Journal of Theoretical Biology 373:62–72.

Haig, D. 2002. Genomic Imprinting and Kinship. Rutgers University Press, New Brunswick, N.J.

Hamilton, W. D. 1963. Evolution of altruistic behavior. The American Naturalist 97:354–356.

Hamilton, W. D. 1964a. The genetical evolution of social behaviour. I. Journal of Theoretical Biology 7:1–16.

Hamilton, W. D. 1964b. The genetical evolution of social behaviour. II. Journal of Theoretical Biology 7:17–52.

Hamilton, W. D. 1970. Selfish and spiteful behaviour in an evolutionary model. Nature 228:1218–1220.

Hamilton, W. D. 1972. Altruism and related phenomena, mainly in social insects. Annual Review of Ecology and Systematics 3:193–232.

Hamilton, W. D. 1975. Innate social aptitudes of man: an approach from evolutionary genetics. Pages 133-155 in R. Fox, editor. Biosocial Anthropology, Wiley, New York.

Hamilton, W. D., and R. M. May. 1977. Dispersal in stable habitats. Nature 269:578–581.

Lande, R., and S. J. Arnold. 1983. The measurment of selection on correlated characters. Evolution 37:1210–1226.

Maynard Smith, J. M., and E. Szathmáry. 1995. The Major Transitions in Evolution. W.H. Freeman Spektrum, Oxford, UK.

Moore, A. J., E. D. Brodie, and J. B. Wolf. 1997. Interacting phenotypes and the evolutionary process. 1. Direct and indirect genetic effects of social interactions. Evolution 51:1352–1362.

Nowak, M. A., A. McAvoy, B. Allen, and E. O. Wilson. 2017. The general form of Hamilton’s rule makes no predictions and cannot be tested empirically. Proceedings of the National Academy of Sciences of the USA 114:5665–5670.

Okasha, S. 2006. Evolution and the Levels of Selection. Oxford University Press, Oxford, UK.

Okasha, S. 2016. On Hamilton’s rule and inclusive fitness theory with nonadditive payoffs. Philosophy of Science 83:873–883.

Price, G. R. 1970. Selection and covariance. Nature 227:520–521.

Price, G. R. 1972. Extension of covariance selection mathematics. Annals of Human Genetics 35:485–490.

Pulliam, H. R. 1988. Sources, sinks, and population regulation. The American Naturalist 132:652–661.

Queller, D. C. 1992a. A general model for kin selection. Evolution 46:376–380.

Queller, D. C. 1992b. Quantitative genetics, inclusive fitness, and group selection. The American Naturalist 139:540–558.

Queller, D. C. 2017. Fundamental theorems of evolution. The American Naturalist 189:345–353.

Rodrigues, A. M. M. 2018. Demography, life history and the evolution of age-dependent social behaviour. Journal of Evolutionary Biology 31:1340–1353.

Rodrigues, A. M. M., and A. Gardner. 2013. Evolution of helping and harming in heterogeneous groups. Evolution 67:2284–2298.

Rousset, F. 2015. Regression, least squares, and the general version of inclusive fitness. Evolution 69:2963–2970.

Taylor, P. D. 1990. Allele-frequency change in a class-structured population. The American Naturalist 135:95–106.

Taylor, P. D., and S. A. Frank. 1996. How to make a kin selection model. Journal of Theoretical Biology 180:27–37.

Trivers, R. L. 1974. Parent-offspring conflict. American Zoologist 14:249–264.

van Veelen, M. 2009. Group selection, kin selection, altruism and cooperation: When inclusive fitness is right and when it can be wrong. Journal of Theoretical Biology 259:589–600.

West, S. A. 2010. Sex Allocation. Princeton University Press, Princeton, NJ.

West, S. A., and A. Gardner. 2013. Adaptation and inclusive fitness. Current Biology 23:R577–R584.

Whiteley, M., S. P. Diggle, and E. P. Greenberg. 2017. Progress in and promise of bacterial quorum sensing research. Nature 551:313–320.

Wilson, D. S., and E. O. Wilson. 2007. Rethinking the theoretical foundation of sociobiology. Quarterly Review of Biology 82:327–348.

