## Appendix for "Using the Price equation to detect inclusive fitness in class-structured populations"

António M. M. Rodrigues[1,*]

1. Department of Ecology & Evolutionary Biology, Yale University, New Haven, 165 Prospect Street, CT 06511, USA.

*

**Table of contents**

A. The Price equation in class-structured populations

A.1. General derivations

A.2. Between-class covariance

A.3. Within-class selection

A.4. The Price equation in class-structured populations

B. The Price equation: Fecundity and Survival effects

B.1. Fecundity effects

B.2. Survival effects

C. Regression analysis:

C.1. Between-class covariance

C.2. Within-class selection: Kin selection

### A. The Price equation in class-structured populations

#### A.1. General derivations

In its general form, the Price equation maps the relationship between two sets of entities (Price 1970, Gardner 2008, Frank 2012). Usually, one set is called the parental population, the other the offspring population, or more generally, the descendant population. Each offspring is connected to a parent ( Gardner 2008, but see Kerr and Godfrey-Smith 2009 ). In addition, each entity is characterised by a breeding value, denoted by *g*, and a reproductive success, denoted by *w*. Given this general set of assumptions, and neglecting transmission biases, the change in average breeding value owing to the action of Natural Selection (NS), denoted by $\Delta_{\mathrm{NS}}\bar{g}$, is given by

$$\Delta_{\mathrm{NS}}\bar{g} = \frac{1}{n}\sum_{\mathrm{i}=1}^{n}\frac{w_{\mathrm{i}}}{\bar{w}}g_{\mathrm{i}} - \frac{1}{n}\sum_{\mathrm{i}=1}^{n}g_{\mathrm{i}}, \tag{A1.1}$$

where $n$ is the number of individuals (or entities) in the population; $w_{\mathrm{i}}$ is the reproductive success of the i*th* individual in the population; $g_{\mathrm{i}}$ is the breeding value of i*th* individual; and $\bar{w}$ is the average reproductive success in the population. The first term in the right-hand side of equation (A1.1) is the average breeding value in the offspring population; the second term is the average breeding value in the parental population. Note that the average reproductive success in the population is given by $\bar{w} = (1/n)\sum_{\mathrm{i}=1}^{n} w_{\mathrm{i}}$, and therefore the total number of offspring produced by the parental population is given by $n\bar{w}$. Further, note that the average breeding value is given by $\bar{g} = (1/n)\sum_{\mathrm{i}=1}^{n} g_{\mathrm{i}}$. This formulation of equation (A1.1) assumes that individuals in the population may differ in breeding value but are otherwise identical. More generally, the entities in a population may differ in many other aspects, in which case we can group them into different "classes". If we assume *N* classes in a population, then Equation (A1.1) becomes

$$\Delta\bar{g} = \frac{1}{n}\left(\sum_{\mathrm{j}=1}^{N}\left(\sum_{\mathrm{i}=1}^{n_{\mathrm{j}}}\frac{w_{\mathrm{ij}}}{\bar{w}}g_{\mathrm{ij}}\right)\right) - \frac{1}{n}\left(\sum_{\mathrm{j}=1}^{N}\left(\sum_{\mathrm{i}=1}^{n_{\mathrm{j}}}g_{\mathrm{ij}}\right)\right), \qquad \text{(A1.2)}$$

where $n_{\mathrm{j}}$ is the number of individuals in the *i*th class, and the subscript *ij* denotes the i*th* individual in class-*j*. We can now re-arrange equation (A1.2), which becomes

$$\begin{aligned}\Delta\bar{g} \; &= \sum_{\mathrm{j}=1}^{N}u_{\mathrm{j}}\frac{1}{\bar{w}}\frac{1}{n_{\mathrm{j}}}\left(\sum_{i=1}^{n_{\mathrm{j}}}w_{\mathrm{ij}}g_{\mathrm{ij}} - \bar{w}_{*\mathrm{j}}\sum_{\mathrm{i}=1}^{n_{\mathrm{j}}}g_{\mathrm{ij}}\right) \\ &+ \sum_{\mathrm{j}=1}^{N}u_{\mathrm{j}}\frac{1}{\bar{w}}\bar{g}_{*\mathrm{j}}\left(\bar{w}_{*\mathrm{j}} - w\right)\end{aligned}, \qquad \text{(A1.3)}$$

where $\bar{w}_{*\mathrm{j}} = \left(1/n_{\mathrm{j}}\right)\sum_{\mathrm{i}=1}^{n_{\mathrm{j}}}w_{\mathrm{ij}}$; $\bar{g}_{*\mathrm{j}} = \left(1/n_{\mathrm{j}}\right)\sum_{\mathrm{i}=1}^{n_{\mathrm{j}}}g_{\mathrm{ij}}$; and $u_{\mathrm{j}} = n_{\mathrm{j}}/n$. While the first summation in the right-hand side of the equation captures the effect of selection operating within classes, the last term in the right-hand side of the equation captures the effect of selection operating between classes and / or class-effects. This will be made clear below.

**A.2. Between-class covariance**

Here, we focus on the between-class covariance term, i.e. the last term in equation (A1.3). We can expand this term, and we obtain

$$\sum_{\mathrm{j}=1}^{N}u_{\mathrm{j}}\frac{1}{\bar{w}}\bar{g}_{*\mathrm{j}}\left(\bar{w}_{*\mathrm{j}} - w\right) = \frac{1}{\bar{w}}\frac{1}{n}\sum_{\mathrm{j}=1}^{N}\sum_{\mathrm{i}=1}^{n_{\mathrm{j}}}\left(g_{\mathrm{ij}} - \bar{g}\right)\left(\bar{w}_{*\mathrm{j}} - \bar{w}\right). \qquad \text{(A2.1)}$$

Note that this is the definition of covariance, and therefore we have

$$\frac{1}{\bar{w}}\frac{1}{n}\sum_{\mathrm{j}=1}^{N}\sum_{\mathrm{i}=1}^{n_{\mathrm{j}}}\left(g_{\mathrm{ij}}-\bar{g}\right)\left(\bar{w}_{*\mathrm{j}}-\bar{w}\right)=\frac{1}{\bar{w}}cov_{\mathrm{B}}\left(\bar{w}_{*\mathrm{j}},g_{\mathrm{ij}}\right). \quad \text{(A2.2)}$$

where the subscript "B" indicates that the covariance is taken between classes and across all individuals in the population.

#### A.3. Within-class selection

Let us now focus on the within-class selection term in the Price equation, i.e. the summation term in the right-hand side of equation (A1.3). If we expand, and given the definition of covariance, we obtain

$$\frac{1}{\bar{w}}\sum_{\mathrm{j}=1}^{N}u_{\mathrm{j}}\frac{1}{n_{\mathrm{j}}}\left(\sum_{\mathrm{i}=1}^{n_{\mathrm{j}}}w_{\mathrm{ij}}g_{\mathrm{ij}}-\bar{w}_{*\mathrm{j}}\sum_{\mathrm{i}=1}^{n_{\mathrm{j}}}g_{\mathrm{ij}}\right)=\frac{1}{\bar{w}}\sum_{\mathrm{j}=1}^{N}u_{\mathrm{j}}cov_{\mathrm{W}}\left(w_{\mathrm{ij}},g_{\mathrm{ij}}\right). \quad \text{(A3.1)}$$

where the subscript "W" indicates that the covariance is taken within each class and across the individuals in that class. When an individual can have *N* types of offspring, its reproductive success is given by

$$w_{\mathrm{ij}}=\sum_{\mathrm{l}=1}^{N}w_{\mathrm{ij}\rightarrow\mathrm{l}}v_{\mathrm{l}}, \quad \text{(A3.2)}$$

where $v_{\mathrm{l}}$ is the reproductive value of each offspring (or future reproductive value). If we put equation (A3.2) into expression (A3.1), we obtain

$$\frac{1}{\bar{w}}\sum_{\mathrm{j}=1}^{N}u_{\mathrm{j}}cov_{\mathrm{W}}\left(w_{\mathrm{ij}},g_{\mathrm{ij}}\right)=\frac{1}{\bar{w}}\sum_{\mathrm{j}=1}^{N}u_{\mathrm{j}}\sum_{\mathrm{l}=1}^{N}v_{\mathrm{l}}cov_{\mathrm{W}}\left(w_{\mathrm{ij}\rightarrow\mathrm{l}},g_{\mathrm{ij}}\right). \quad \text{(A3.3)}$$

#### A.4. The Price equation for class-structured populations

There are two useful forms of the Price equation for class-structured populations. One comes from considering the expressions (A2.2) and (A3.1), in which case equation (A1.3) becomes

$$\begin{aligned}\Delta\bar{g} &= \tfrac{1}{\bar{w}}\textstyle\sum_{\mathrm{j}=1}^{N} u_{\mathrm{j}} cov_{\mathrm{W}}\left(w_{\mathrm{ij}}, g_{\mathrm{ij}}\right) \\ &+ \tfrac{1}{\bar{w}} cov_{\mathrm{B}}\left(\bar{w}_{*\mathrm{j}}, g_{\mathrm{ij}}\right)\end{aligned} . \quad \text{(A4.1)}$$

Another useful form of the Price equation for class-structured populations comes from expressions (A2.2) and expressions (A3.1), in which case equation (A1.3) becomes

$$\begin{aligned}\Delta\bar{g} &= \tfrac{1}{\bar{w}}\textstyle\sum_{\mathrm{j}=1}^{N} u_{\mathrm{j}} \sum_{\mathrm{l}=1}^{N} v_{\mathrm{l}} cov_{\mathrm{W}}\left(w_{\mathrm{ij}\to\mathrm{l}}, g_{\mathrm{ij}}\right) \\ &+ \tfrac{1}{\bar{w}} cov_{\mathrm{B}}\left(\bar{w}_{*\mathrm{j}}, g_{\mathrm{ij}}\right)\end{aligned} . \quad \text{(A4.2)}$$

This gives an exact description of change in mean breeding value due to the action of natural selection and class-effects in class-structured populations.

### B. The Price equation: Fecundity and Survival effects

#### B.1. Fecundity effects

Let us assume that there is standing genetic variation for fecundity only. Consider the fitness of the *i*th individual in class *k*, given by its contribution to the different classes of individuals in the populations. Thus, we have

$$w_{\mathrm{ik}} = \sum_{\mathrm{l}=1}^{\mathrm{N}} w_{\mathrm{ik}\to\mathrm{l}}. \qquad \text{(B1.1)}$$

We can partition the fitness of the focal individual into class-specific fecundity, denoted by $f_{\mathrm{ik}\to\mathrm{l}}$, and reproductive value of offspring, denoted by $V_{\mathrm{l}}$. This is given by

$$w_{\mathrm{ik}} = \sum_{\mathrm{l}=1}^{\mathrm{N}} f_{\mathrm{ik}\to\mathrm{l}} V_{\mathrm{l}}. \qquad \text{(B1.2)}$$

The class-specific fertility is given by the overall fecundity, denoted by $f_{\mathrm{ik}}$, times the fraction of offspring that belong to each class, which is given by $q_{\mathrm{k}\to\mathrm{l}}$. Hence, we have

$$f_{\mathrm{ik}\to\mathrm{l}} = f_{\mathrm{ik}} q_{\mathrm{k}\to\mathrm{l}}. \qquad \text{(B1.3)}$$

We can now put equation (B1.3) into equation (B1.2) to obtain

$$w_{\mathrm{ik}} = \sum_{\mathrm{l}=1}^{\mathrm{N}} f_{\mathrm{ik}\to\mathrm{l}} V_{\mathrm{l}} = f_{\mathrm{ik}} \sum_{\mathrm{l}=1}^{\mathrm{N}} q_{\mathrm{k}\to\mathrm{l}} V_{\mathrm{l}}. \qquad \text{(B1.4)}$$

and then calculate the average fitness of class *k*, which is given by

$$\bar{w}_{*\mathrm{k}} = \frac{1}{n_{\mathrm{k}}} \sum_{\mathrm{i}=1}^{n_{\mathrm{k}}} w_{\mathrm{ik}} = \frac{1}{n_{\mathrm{k}}} \sum_{\mathrm{i}=1}^{n_{\mathrm{k}}} \sum_{\mathrm{l}=1}^{\mathrm{N}} w_{\mathrm{ik}\to\mathrm{l}} = \frac{1}{n_{\mathrm{k}}} \sum_{\mathrm{i}=1}^{n_{\mathrm{k}}} f_{\mathrm{ik}} \sum_{\mathrm{l}=1}^{\mathrm{N}} q_{\mathrm{k}\to\mathrm{l}} V_{\mathrm{l}}. \qquad \text{(B1.5)}$$

which allows us to define the expected reproductive value of offspring as

$$\bar{V} = \sum_{\mathrm{l=1}}^{\mathrm{N}} q_{\mathrm{k\to l}} V_{\mathrm{l}}. \quad \text{(B1.6)}$$

Equation (B1.4) and equation (B1.5) can now be written as

$$w_{\mathrm{ik}} = f_{\mathrm{ik}} \bar{V}, \text{ and} \quad \text{(B1.7)}$$

$$\bar{w}_{*\mathrm{j}} = \left(\frac{1}{n_{\mathrm{j}}} \sum_{\mathrm{i=1}}^{n_{\mathrm{j}}} f_{\mathrm{ij}}\right) \bar{V} = \bar{f}_{*\mathrm{j}} \bar{V}, \quad \text{(B1.8)}$$

respectively, where:

$$\bar{f}_{*\mathrm{j}} = \frac{1}{n_{\mathrm{j}}} \sum_{\mathrm{i=1}}^{n_{\mathrm{j}}} f_{\mathrm{ij}}. \quad \text{(B1.9)}$$

Let us now assume that mother vary in fecundity but that offspring are produced in exactly the same proportion by all mothers in the population. Then, the reproductive value of a focal mother can be written as

$$v_{\mathrm{ik\to l}} = f_{\mathrm{ik}} q_{\mathrm{l}} V_{\mathrm{l}}. \quad \text{(B1.10)}$$

We can now put equations (B1.10) and (B1.8) into the Price equation, given by equation (B4.1), to obtain

$$\bar{w}\Delta\bar{g} = \sum_{\mathrm{j=1}}^{\mathrm{N}} u_{\mathrm{j}} \sum_{\mathrm{l=1}}^{N} cov_{\mathrm{W}}\left(f_{\mathrm{ik}} q_{\mathrm{l}} V_{\mathrm{l}}, g_{\mathrm{ij}}\right) + cov_{\mathrm{B}}\left(\bar{f}_{*\mathrm{j}} \bar{V}, g_{\mathrm{ij}}\right). \quad \text{(B1.11)}$$

If we expand equation (B1.11), we get

$$\bar{w}\Delta\bar{g} = \sum_{\mathrm{j}=1}^{\mathrm{N}} u_{\mathrm{j}} cov_{\mathrm{W}}\left(f_{\mathrm{ik}}, g_{\mathrm{ij}}\right) \sum_{\mathrm{l}=1}^{N} q_{\mathrm{l}} V_{\mathrm{l}} + cov_{\mathrm{B}}\left(\bar{f}_{\mathrm{ij}}, g_{\mathrm{ij}}\right)\bar{V}. \quad \text{(B1.12)}$$

Given equation (B1.6), we can now write:

$$\bar{w}\Delta\bar{g} = \sum_{\mathrm{j}=1}^{\mathrm{N}} u_{\mathrm{j}} cov_{\mathrm{W}}\left(f_{\mathrm{ik}}, g_{\mathrm{ij}}\right)\bar{V} + cov_{\mathrm{B}}\left(\bar{f}_{*\mathrm{j}}, g_{\mathrm{ij}}\right)\bar{V}. \quad \text{(B1.13)}$$

Equation (B1.13) shows that when the standing genetic variation affects maternal fertility selection within classes depends on the covariance between breeding value and fertility, while selection between classes and / or class-effects depends on the covariance between breeding value and average class fertility.

**B.2. Survival effects**

Let us now assume that there is standing genetic variation for survival only. As above, consider the fitness of the *i*th individual in class *k*, given by its contribution to the different classes of individuals in the population. Thus, we have

$$w_{\mathrm{ik}} = \sum_{\mathrm{l}=1}^{\mathrm{N}} w_{\mathrm{ik}\to\mathrm{l}}. \quad \text{(B2.1)}$$

We can partition the class-specific fitness of a focal individual into survival, denoted by $s_{\mathrm{ik}\to\mathrm{l}}$, and future reproductive value, denoted by $v_{\mathrm{l}}$, as follows:

$$w_{\mathrm{ik}\to\mathrm{l}} = s_{\mathrm{ik}\to\mathrm{l}} v_{\mathrm{l}}. \quad \text{(B2.2)}$$

Thus, equation (B2.1) becomes

$$w_{\mathrm{ik}} = \sum_{\mathrm{l}=1}^{\mathrm{N}} s_{\mathrm{ik}\to\mathrm{l}} v_{\mathrm{l}}. \tag{B2.3}$$

The class-specific survival is given by the overall survival, denoted by $s_{\mathrm{ik}}$, times the probability of becoming a class-*l* individual, which is given by $q_{\mathrm{k}\to\mathrm{l}}$. For instance, this may denote the probability that an age-4 individual becomes an age-5 individual. Hence, we have

$$s_{\mathrm{ik}\to\mathrm{l}} = s_{\mathrm{ik}} q_{\mathrm{k}\to\mathrm{l}}. \tag{B2.4}$$

We can now put equation (B2.4) into the equation (B2.3) to obtain

$$w_{\mathrm{ik}} = \sum_{\mathrm{l}=1}^{\mathrm{N}} s_{\mathrm{ik}\to\mathrm{l}} v_{\mathrm{l}} = s_{\mathrm{ik}} \sum_{\mathrm{l}=1}^{\mathrm{N}} q_{\mathrm{k}\to\mathrm{l}} v_{\mathrm{l}}. \tag{B2.5}$$

We can now calculate the average fitness of class *k* as

$$\bar{w}_{*\mathrm{k}} = \frac{1}{n_{\mathrm{k}}} \sum_{\mathrm{i}=1}^{n_{\mathrm{k}}} w_{\mathrm{ik}} = \frac{1}{n_{\mathrm{k}}} \sum_{\mathrm{i}=1}^{n_{\mathrm{k}}} \sum_{\mathrm{l}=1}^{\mathrm{N}} w_{\mathrm{ik}\to\mathrm{l}} = \frac{1}{n_{\mathrm{k}}} \sum_{\mathrm{i}=1}^{n_{\mathrm{k}}} s_{\mathrm{ik}} \sum_{\mathrm{l}=1}^{\mathrm{N}} q_{\mathrm{k}\to\mathrm{l}} v_{\mathrm{l}}. \tag{B2.6}$$

and define the expected reproductive value of an adult as

$$\bar{v}_{\mathrm{k}} = \sum_{\mathrm{l}=1}^{\mathrm{N}} q_{\mathrm{k}\to\mathrm{l}} v_{\mathrm{l}}. \tag{B2.7}$$

Equation (B2.5) and equation (B2.6) can now be written as

$$w_{\mathrm{ik}} = s_{\mathrm{ik}}\bar{v}_{\mathrm{k}}, \text{ and} \tag{B2.8}$$

$$\bar{w}_{*\mathrm{k}} = \left(\frac{1}{n_{\mathrm{k}}}\sum_{\mathrm{i=1}}^{n_{\mathrm{k}}} s_{\mathrm{ik}}\right)\bar{v}_{\mathrm{k}} = \bar{s}_{*\mathrm{k}}\bar{v}_{\mathrm{k}}, \tag{B2.9}$$

respectively, where:

$$\bar{s}_{*\mathrm{k}} = \frac{1}{n_{\mathrm{k}}}\sum_{\mathrm{i=1}}^{n_{\mathrm{k}}} s_{\mathrm{ik}}. \tag{B2.10}$$

We can now put equations (B2.8) and (B2.10) into the Price equation (Eqn. A4.1), to obtain

$$\bar{w}\Delta\bar{g} = \left(\sum_{\mathrm{j=1}}^{\mathrm{N}} u_{\mathrm{j}} \sum_{\mathrm{l=1}}^{N} cov_{\mathrm{W}}\left(s_{\mathrm{ij}}q_{\mathrm{j\to l}}v_{\mathrm{l}}, g_{\mathrm{ij}}\right)\right) + cov_{\mathrm{B}}\left(\bar{s}_{*\mathrm{j}}\bar{v}_{\mathrm{j}}, g_{\mathrm{ij}}\right). \tag{B2.11}$$

If we re-arrange

$$\bar{w}\Delta\bar{g} = \left(\sum_{\mathrm{j=1}}^{\mathrm{N}} u_{\mathrm{j}} cov_{\mathrm{W}}\left(s_{\mathrm{ij}}, g_{\mathrm{ij}}\right) \sum_{\mathrm{l=1}}^{N} q_{\mathrm{j\to l}}v_{\mathrm{l}}\right) + cov_{\mathrm{B}}\left(\bar{s}_{*\mathrm{j}}\bar{v}_{\mathrm{j}}, g_{\mathrm{ij}}\right). \tag{B2.12}$$

From equation (B2.7), we finally obtain

$$\bar{w}\Delta\bar{g} = \left(\sum_{\mathrm{j=1}}^{\mathrm{N}} u_{\mathrm{j}} cov_{\mathrm{W}}\left(s_{\mathrm{ij}}, g_{\mathrm{ij}}\right)\bar{v}_{\mathrm{j}}\right) + cov_{\mathrm{B}}\left(\bar{s}_{*\mathrm{j}}\bar{v}_{\mathrm{j}}, g_{\mathrm{ij}}\right). \tag{B2.13}$$

Equation (B2.13) shows that selection within classes depends on the covariance between breeding value and survival whereas selection between classes and / or class effects depends on the covariance between breeding value and average class survival.

### C. Regression analysis

#### C.1. Between-class covariance

We can define a statistical model to describe the average fitness of a focal individual. First, we consider the class phenotype, which is given by

$$\sigma_{\mathrm{ij}} = \frac{\left(\bar{w}_{*\mathrm{j}} - \min(\bar{w}_{*\mathrm{j}})\right)}{\max\left(\bar{w}_{*\mathrm{j}} - \min(\bar{w}_{*\mathrm{j}})\right)}, \tag{C1.1}$$

where the function min() returns the lowest value and the function max() returns the largest value. Notice that the phenotype is proportional to the average fitness in the class. Thus, a class where the mean fitness is lower will have a lower number while a class where the mean fitness is higher will have a larger number. Further, we normalise this quantity such that it is bounded between 0 and 1. The class with the highest mean fitness will be assigned a number 1. To visualise this, imagine that the mean fitness of individuals depends on their size, and the association between size and fitness is perfect. The statistical model that predicts mean fitness is then given by

$$\bar{w}_{*\mathrm{j}} = \beta_{\mathrm{c0}} + \beta_{\mathrm{c}}\sigma_{\mathrm{ij}} + \varepsilon_{\mathrm{ij}} \quad i \in (1, \dots, n_{\mathrm{j}}), \tag{C1.2}$$

where $\beta_{\mathrm{c0}}$ is the intercept of the regression analysis, and $\beta_{\mathrm{c}}$ is the effect of class phenotype on average class fitness. Now, remember that the between-class covariance term in equation (A4.1) is given by

$$cov_{\mathrm{B}}\left(\bar{w}_{*\mathrm{j}}, g_{\mathrm{ij}}\right). \tag{C1.3}$$

If we now put the statistical model (C1.2) into the covariance term (C1.3), we obtain

$$cov_{\mathrm{B}}\left(\bar{w}_{*\mathrm{j}}, g_{\mathrm{ij}}\right) = cov_{\mathrm{B}}\left(\beta_{\mathrm{c}0} + \beta_{\mathrm{c}}\sigma_{\mathrm{ij}} + \varepsilon_{\mathrm{ij}}, g_{\mathrm{ij}}\right). \tag{C1.4}$$

It is important to bear in mind that the covariance is calculated across all individuals in the population. We can now expand equation (C1.4), such that we obtain:

$$cov_{\mathrm{B}}\left(\beta_{\mathrm{c}0} + \beta_{\mathrm{c}}\sigma_{\mathrm{ij}} + \varepsilon_{\mathrm{ij}}, g_{\mathrm{ij}}\right) = \beta_{\mathrm{c}} cov_{\mathrm{B}}\left(\sigma_{\mathrm{ij}}, g_{\mathrm{ij}}\right) = \beta_{\mathrm{c}} \frac{cov_{\mathrm{B}}\left(\sigma_{\mathrm{ij}}, g_{\mathrm{ij}}\right)}{var_{\mathrm{B}}\left(g_{\mathrm{ij}}\right)} var_{\mathrm{B}}\left(g_{\mathrm{ij}}\right). \tag{C1.5}$$

Finally, let us defined $d_{\mathrm{c}} = \beta_{\mathrm{c}}$ and $r_{\mathrm{c}} = cov_{\mathrm{B}}\left(\sigma_{\mathrm{ij}}, g_{\mathrm{ij}}\right) / var_{\mathrm{B}}\left(g_{\mathrm{ij}}\right)$ and we finally get

$$cov_{\mathrm{B}}\left(\bar{w}_{*\mathrm{j}}, g_{\mathrm{ij}}\right) = d_{\mathrm{c}} r_{\mathrm{c}} var_{\mathrm{B}}\left(g_{\mathrm{ij}}\right). \tag{C1.6}$$

Thus, the covariance between breeding value and average class fitness is given by the benefits of class membership, $d_{\mathrm{c}}$, the correlation between class membership and breeding value, i.e. $r_{\mathrm{c}}$, and the amount of additive genetic variation in the population, i.e. $var_{\mathrm{B}}\left(g_{\mathrm{ij}}\right)$.

**C.2. Selection within classes: Kin selection**

Now consider a statistical model of the fecundity of individuals in each one of the population classes. First, let $\beta_{0\mathrm{j}\to\mathrm{l}}$ be the intercept of the statistical model that describes the fecundity of parents in class-*j* through offspring that become class-*l* offspring. Second, let $\beta_{\mathrm{ij}\to\mathrm{l}}$ be the partial regression coefficient that gives the effect of the focal individual's breeding value on its own fecundity through offspring that become class-*l* individuals. Third, let $\beta_{\mathrm{i}\sigma\to\mathrm{j}\to\mathrm{l}}$ be the partial regression coefficient that gives the effect of a class-$\sigma$ social partners on the fitness of class-*j* individuals through their offspring that become class-*l* individuals. Fourth, let the actors of the behaviour belong to class σ and let the set Ω be the set of all classes that contain the actors of the behaviour. Given this statistical model of the fitness of individuals, the Price equation becomes

$$\Delta\bar{g} = \frac{1}{\bar{w}}\left(\sum_{\mathrm{j}=1}^{\mathrm{N}} u_{\mathrm{j}} \sum_{\mathrm{l}=1}^{\mathrm{N}} v_{\mathrm{j}} cov_{\mathrm{W}}\left(\beta_{\mathrm{ij}\to\mathrm{l}} g_{\mathrm{ij}} + \sum_{\sigma\in\Omega} \beta_{\mathrm{i}\sigma\to\mathrm{j}\to\mathrm{l}} G_{\mathrm{i}\sigma}, g_{\mathrm{ij}}\right)\right) + \frac{1}{\bar{w}} d_{\mathrm{c}} r_{\mathrm{c}} var_{\mathrm{B}}\left(g_{\mathrm{ij}}\right). \qquad \text{(C2.1)}$$

where $G_{\mathrm{i}\sigma}$ is the breeding value of the class-σ that are the social partners of the class-*j* individuals. Given the additive property of the covariance function, the covariance of a sum becomes the sum of covariances as follows:

$$\begin{aligned}\Delta\bar{g} &= \frac{1}{\bar{w}}\left(\sum_{\mathrm{j}=1}^{\mathrm{N}} u_{\mathrm{j}} \sum_{\mathrm{l}=1}^{\mathrm{N}} v_{\mathrm{j}} \left(cov_{\mathrm{W}}\left(\beta_{\mathrm{ij}\to\mathrm{l}} g_{\mathrm{ij}}, g_{\mathrm{ij}}\right) + cov_{\mathrm{W}}\left(\sum_{\sigma\in\Omega} \beta_{\mathrm{i}\sigma\to\mathrm{j}\to\mathrm{l}} G_{\mathrm{i}\sigma}, g_{\mathrm{ij}}\right)\right)\right) \\ &\quad + \frac{1}{\bar{w}} d_{\mathrm{c}} r_{\mathrm{c}} var_{\mathrm{B}}\left(g_{\mathrm{ij}}\right)\end{aligned}. \qquad \text{(C2.2)}$$

We can now expand the first term in the right-hand side to obtain

$$\begin{aligned}\bar{w}\Delta\bar{g} &= \sum_{\mathrm{j}=1}^{\mathrm{N}} u_{\mathrm{j}} \sum_{\mathrm{l}=1}^{\mathrm{N}} v_{\mathrm{j}} \beta_{\mathrm{ij}\to\mathrm{l}} cov_{\mathrm{W}}\left(g_{\mathrm{ij}}, g_{\mathrm{ij}}\right) \\ &\quad + \sum_{\mathrm{j}=1}^{\mathrm{N}} u_{\mathrm{j}} \sum_{\mathrm{l}=1}^{\mathrm{N}} v_{\mathrm{j}} \sum_{\sigma\in\Omega} \beta_{\mathrm{i}\sigma\to\mathrm{j}\to\mathrm{l}} cov_{\mathrm{W}}\left(G_{\mathrm{i}\sigma}, g_{\mathrm{ij}}\right) + d_{\mathrm{c}} r_{\mathrm{c}} var_{\mathrm{B}}\left(g_{\mathrm{ij}}\right)\end{aligned}. \qquad \text{(C2.3)}$$

Let us assume that mothers produce the same proportion of offspring types, and therefore the effects of social partners only affect the fecundity of a focal individual. Then $\beta_{\mathrm{ij}\to\mathrm{l}} = \beta_{\mathrm{ij}} V_{\mathrm{l}}$ and

$\beta_{i\sigma\to j\to l} = \beta_{i\sigma\to j} V_l$, where I use capital "V" to highlight that the reproductive values are offspring reproductive values. We can then write:

$$\bar{w}\Delta\bar{g} = \sum_{j=1}^{N} u_j \sum_{l=1}^{N} \beta_{ij} cov_W\left(g_{ij}, g_{ij}\right) V_l + \sum_{j=1}^{N} u_j \sum_{l=1}^{N} \sum_{\sigma\in\Omega} \beta_{i\sigma\to j} cov_W\left(G_{i\sigma}, g_{ij}\right) V_l + \beta_c r_c var_B\left(g_{ij}\right). \quad \text{(C2.4)}$$

Now assume, without loss of generality, that the behaviour is only performed by individuals in class $\alpha$; thus, $\beta_{ij} = 0$ when $ij \neq \alpha$. Further, this means that the class of actors is class $\alpha$ only, which means that $\Omega = \alpha$, and $\sum_{\sigma\in\Omega} \beta_{i\sigma\to j} cov_W\left(G_{i\sigma}, g_{ij}\right) V_l = \beta_{\alpha\to j} cov_W\left(G_{i\alpha}, g_{ij}\right) V_l$, which implies that equation (C2.4) becomes

$$\bar{w}\Delta\bar{g} = \sum_{j=1}^{N} u_j \sum_{l=1}^{N} \beta_{ij} cov_W\left(g_{ij}, g_{ij}\right) V_l + \sum_{j=1}^{N} u_j \sum_{l=1}^{N} \beta_{i\sigma\to j} cov_W\left(G_{i\sigma}, g_{ij}\right) V_l + d_c r_c var_B\left(g_{ij}\right). \quad \text{(C2.5)}$$

Now let class α individuals only provide help to a sub-set of individuals, and let Θ represent the set of all individuals that receive benefits from individuals in class α.

$$\bar{w}\Delta\bar{g} = u_\alpha \sum_{l=1}^{N} \beta_{i\alpha} cov_W(g_{i\alpha}, g_{i\alpha}) V_l + \sum_{\rho=\Theta} u_\rho \sum_{l=1}^{N} \beta_{i\alpha\to\rho} cov_W\left(G_{i\alpha}, g_\rho\right) V_l + d_c r_c var_B\left(g_{ij}\right). \quad \text{(C2.6)}$$

Note that the partial regression coefficient gives the effect of the actor on the fecundity of the recipients, and the covariances are between the actor and the recipients. Neither of these two quantities depend on the class of offspring, and therefore we can write equation (C2.6) as follows:

$$\bar{w}\Delta\bar{g} = \begin{array}{l} u_\alpha \beta_{i\alpha} cov_W(g_{i\alpha}, g_{i\alpha}) \sum_{l=1}^{N} q_l V_l + \sum_{\rho=\Theta} u_\rho \beta_{\alpha\to\rho} cov_W(G_{i\alpha}, g_\rho) \sum_{l=1}^{N} q_l V_l \\ +d_c r_c var_B(g_{ij}) \end{array}. \quad \text{(C2.7)}$$

Now notice that $\sum_{l=1}^{N} q_l V_l$ is the expected reproductive value of an offspring, which we can denote by $\bar{V} = \sum_{l=1}^{N} q_l V_l$. With this notation, equation (C2.7) becomes

$$\bar{w}\Delta\bar{g} = \begin{array}{l} u_\alpha \beta_{i\alpha} cov_W(g_{i\alpha}, g_{i\alpha}) \bar{V} + \sum_{\rho=\Theta} u_\rho \beta_{i\alpha\to\rho} cov_W(g_{i\alpha}, g_{i\rho}) \bar{V} \\ +d_c r_c var_B(g_{ij}) \end{array}. \quad \text{(C2.8)}$$

Now notice that because $cov_W(g_{i\alpha}, g_{i\alpha}) = var_W(g_{i\alpha})$ and $cov_W(g_{i\alpha}, g_{i\rho}) = cov_W(g_{i\rho}, g_{i\alpha})$, equation (C2.8) becomes

$$\bar{w}\Delta\bar{g} = \begin{array}{l} u_\alpha \beta_{i\alpha} var_W(g_{i\alpha}) \bar{V} + \sum_{\rho=\Theta} u_\rho \beta_{i\alpha\to\rho} cov_W(g_{i\rho}, g_{i\alpha}) \bar{V} \\ +d_c r_c var_B(g_{ij}) \end{array}. \quad \text{(C2.9)}$$

We can now expand equation (C2.9) to get

$$\bar{w}\Delta\bar{g} = \begin{array}{l} u_\alpha \left(\beta_{i\alpha} + \sum_{\rho=\Theta} \frac{u_\rho}{u_\alpha} \beta_{i\alpha\to\rho} \frac{cov_W(g_{i\rho}, g_{i\alpha})}{var_W(g_{i\alpha})}\right) var_W(g_{i\alpha}) \bar{V} \\ +d_c r_c var_B(g_{ij}) \end{array}. \quad \text{(C2.10)}$$

Notice that $\beta_{i\alpha\to\rho}$ gives the effect of class α individuals on class ρ individuals. Thus, the effect of each class α on each of the class ρ individuals is given by $u_\rho/u_\alpha\, \beta_{i\alpha\to\rho}$. Notice also, that the coefficients of relatedness are given by

$$r_{\alpha\rho} = \frac{cov_W(g_{i\rho}, g_{i\alpha})}{var_W(g_{i\alpha})}. \quad \text{(C2.11)}$$

Equation (C2.10) allows us to recover Hamilton's rule in its traditional form.

$$\bar{w}\Delta\bar{g} = \begin{array}{l} u_\alpha \left(\beta_{i\alpha} + \sum_{\rho=\Theta} \frac{u_\rho \beta_{i\rho\to\alpha}}{u_\alpha} \frac{cov_W(g_{i\rho}, g_{i\alpha})}{var_W(g_{i\alpha})}\right) var_W(g_{i\alpha}) \sum_{l=1}^{N} V_l \\ +d_c r_c var_B(g_{ij}) \end{array}. \qquad \text{(C2.12)}$$

In particular, we find that: $\beta_{i\alpha}$ represents the cost of the behaviour, usually denoted by – *c*; $u_\rho \beta_{i\rho\to\alpha}/u_\alpha$ is the benefit provided by an individual in class α to an individual in class ρ, usually denoted by $b_{\alpha\to\rho}$; and $cov_W(g_{i\rho}, g_{i\alpha})/var_W(g_{i\alpha})$ is the coefficient of relatedness between individuals in class α and individuals in class ρ. In this section we considered the regression method assuming fecundity effects. A similar approach can be taken when one assumes survival effects.

### References


Frank, S. A. 2012. Natural selection. IV. The Price equation. Journal of Evolutionary Biology **25**:1002-1019.

Gardner, A. 2008. The Price equation. Current Biology **18**:R198-R202.

Kerr, B., and P. Godfrey-Smith. 2009. Generalization of the Price equation for evolutionary change. Evolution **63**:531-536.

Price, G. R. 1970. Selection and covariance. Nature **227**:520-521.
